# Analysis of a Mad2 homolog in *Trypanosoma brucei* provides possible hints on the origin of the spindle checkpoint

**DOI:** 10.1101/2020.12.29.424754

**Authors:** Bungo Akiyoshi

## Abstract

The spindle checkpoint is a surveillance mechanism that ensures accurate nuclear DNA segregation in eukaryotes. It does so by delaying the onset of anaphase until all kinetochores have established proper attachments to spindle microtubules. The evolutionary origin of the spindle checkpoint remains unclear. The flagellated kinetoplastid parasite *Trypanosoma brucei* has a nucleus that contains the nuclear genome and a kinetoplast that contains the mitochondrial genome. The kinetoplast is physically linked to the basal body of the flagellum and its segregation is driven by the movement of basal bodies. While there is no strong evidence that *T. brucei* possesses a functional spindle checkpoint, it has been suggested that initiation of cytokinesis may be linked to the completion of kinetoplast segregation or basal body separation in this organism. Interestingly, the only identifiable spindle checkpoint component TbMad2 localizes at the basal body area. Here, I report identification of proteins that co-purified with TbMad2. One protein, which I propose to call TbMBP65, localizes at the basal body area and has a putative Mad2-interacting motif. Interestingly, 26S proteasome subunits also co-purified with TbMad2, raising a possibility that TbMad2 might regulate proteasome activity to regulate or monitor the segregation of basal bodies. I speculate that such a function might represent a prototype of the spindle checkpoint. I also show that TbAUK3, one of the three Aurora kinase homologs in *T. brucei*, localizes at the basal body area from late G1 until the time of kinetoplast separation. Immunoprecipitation of TbAUK3 identified an uncharacterized protein (termed TbABP79) that has a similar localization pattern as TbAUK3. These findings provide a starting point to reveal the function of TbMad2 and TbAUK3 as well as the origin of the spindle checkpoint.

## Introduction

It is essential that cells transmit their genetic material and organelles accurately at each cell division. Kinetochores are the macromolecular protein complexes that assemble onto centromeric DNA and interact with spindle microtubules in eukaryotes (Musacchio and Desai, 2017). Accurate chromosome segregation requires that kinetochores attach to microtubules emanating from opposite poles. The spindle checkpoint is a surveillance system that ensures high fidelity segregation of nuclear DNA (Murray, 2011). In response to defects in kinetochore-microtubule attachments, the spindle checkpoint delays the onset of anaphase by inhibiting the multi-subunit ubiquitin ligase called the anaphase-promoting complex/cyclosome (APC/C) (Musacchio and Salmon, 2007; London and Biggins, 2014; Musacchio, 2015; Alfieri et al., 2017). Once all chromosomes have achieved proper attachments, the spindle checkpoint is satisfied, leading to the ubiquitylation of two key targets (cyclin B and securin) by the APC/C^Cdc20^. Ubiquitylated proteins are subsequently destroyed by the 26S proteasome, triggering chromosome segregation and the exit of mitosis. Spindle checkpoint components include Mad1, Mad2, Mad3 (BubR1), Mps1, Bub1 and Bub3. These proteins are widely conserved among eukaryotes and it is therefore thought that the last eukaryotic common ancestor had a spindle checkpoint (Vleugel et al., 2012; van Hooff et al., 2017; Tromer et al., 2019; Kops et al., 2020). A key player Mad2 is recruited to the kinetochore by direct interaction with Mad1, which promotes conformational conversion of Mad2 so that it that can interact with Cdc20 (De Antoni et al., 2005). Mad1 and Cdc20 have similar Mad2-interacting motifs (K/R-Φ-Φ-x-x-x-x-x-P, where Φ is an aliphatic residue and x is any residue) (Luo et al., 2002; Sironi et al., 2002). Although there have been extensive studies in select model eukaryotes that all belong to Opisthokonta (e.g. yeasts, worms, flies, and human), very little is known about the spindle checkpoint or cell cycle regulation in evolutionarily divergent eukaryotes (Kaczanowski et al., 1985; Morrissette and Sibley, 2002; Vicente and Cande, 2014; Komaki and Schnittger, 2016; Markova et al., 2016).

While segregation of nuclear DNA depends on spindle microtubules in all studied eukaryotes, the mechanism for mitochondrial DNA transmission varies among eukaryotes. Animals have a high copy number of mitochondria and their transmission is thought to occur randomly (Westermann, 2010). In contrast, a single mitochondrion is present in some unicellular eukaryotes, such as *Trypanosoma brucei* (the kinetoplastid parasite that causes human African trypanosomiasis, Discobids), *Plasmodium falciparum* (the parasite that causes malaria, Alveolates), and *Cyanidioschyzon merolae* (red algae, Archaeplastids) (Vickerman, 1962; Itoh et al., 1997; Okamoto et al., 2009; Walker et al., 2011; Keeling and Burki, 2019). *T. brucei* has a kinetoplast that contains mitochondrial DNA (kDNA) and a nucleus that contains nuclear DNA (Figure 1). Kinetoplasts segregate prior to the nuclear division, followed by cytokinesis (Woodward and Gull, 1990; Siegel et al., 2008). It is essential that cells duplicate and segregate these single-copy organelles accurately at each cell division cycle. The kDNA is physically attached to the basal body of the flagellum via the tripartite attachment complex, and its segregation is driven by the separation of basal bodies/flagella (Robinson and Gull, 1991; Ogbadoyi et al., 2003; Schneider and Ochsenreiter, 2018). *T. brucei* does not break down its nuclear envelope (closed mitosis), and an intranuclear mitotic spindle forms in the nucleus during mitosis (Vickerman and Preston, 1970; Ogbadoyi et al., 2000). It is thought that *T. brucei* does not have a canonical spindle checkpoint because cells are unable to halt their cell cycle in response to spindle defects and undergo cytokinesis without a noticeable delay despite a lack of nuclear division (Robinson et al., 1995; Ploubidou et al., 1999; Hayashi and Akiyoshi, 2018), which is consistent with the apparent absence of all the spindle checkpoint components mentioned above (except for a Mad2 homolog called TbMad2, see below) as well as the absence of the well-conserved Mad2-interacting motif in TbCdc20 (Akiyoshi and Gull, 2013). Cyclin B (called CYC6 in *T. brucei*) localizes at the basal body area in S phase and appears in the nucleus in G2 phase until it disappears at the onset of anaphase (Hayashi and Akiyoshi, 2018). Forced stabilization of cyclin B in the nucleus causes the nucleus to arrest in a metaphase-like state without preventing cytokinesis, suggesting that trypanosomes have an ability to regulate the timing of nuclear division by modulating the cyclin B protein level without a canonical spindle checkpoint.

**Figure 1.**
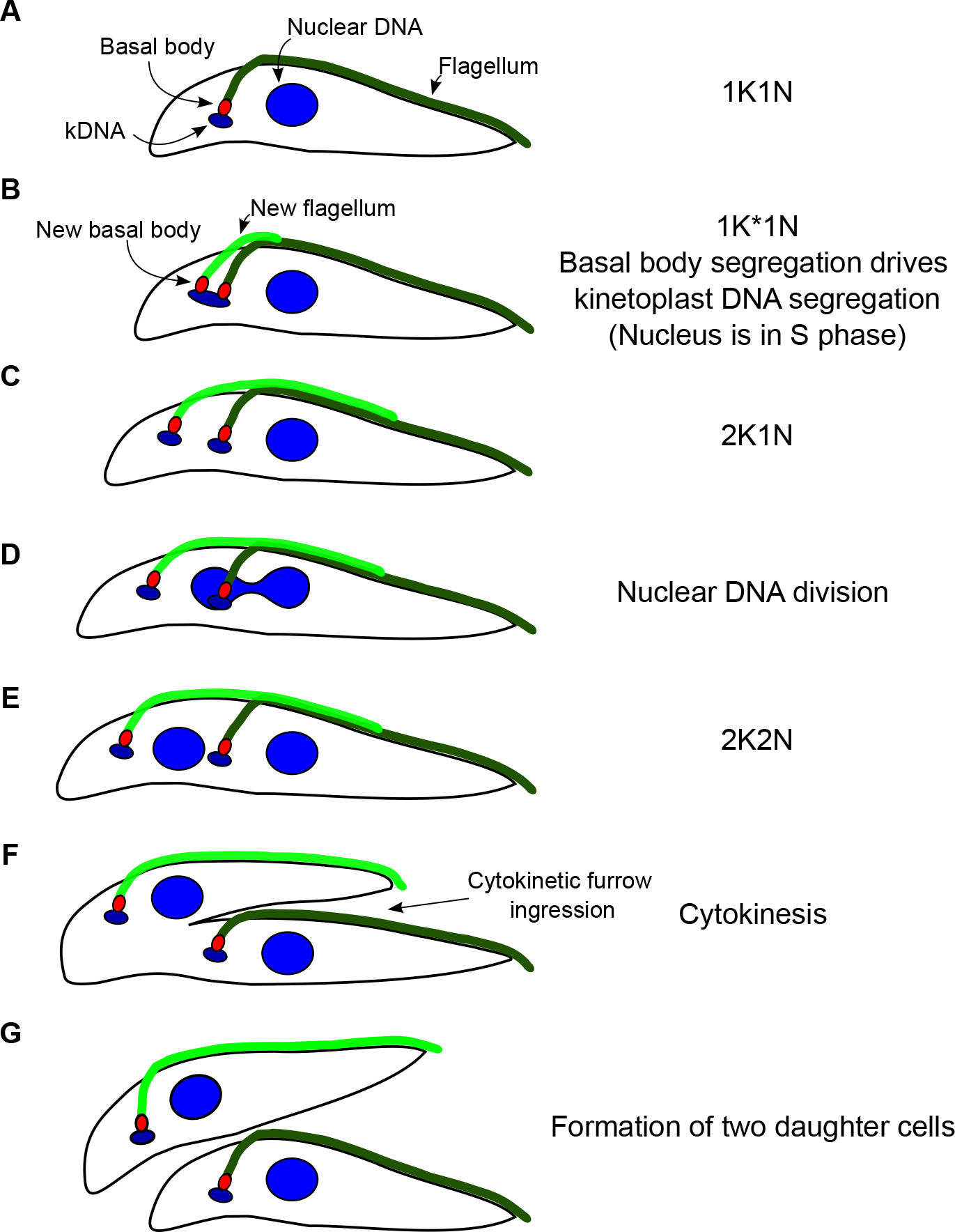
Diagram of the cell division cycle in *Trypanosoma brucei* procyclic (insect) form cells. (A) G1 cells possess a single kinetoplast and nucleus (termed 1K1N) as well as an attached flagellum. (B) As the cell cycle progresses, a new basal body forms and nucleates a new flagellum. The nucleus is still in S phase when kinetoplast DNA (kDNA) has an elongated morphology (termed 1K*1N). (C) Segregation of basal bodies leads to the separation of attached kinetoplast (termed 2K1N). (D) Cells enter nuclear M phase, and chromosome segregation occurs. (E) Nuclear division is complete (termed 2K2N). (F) Cleavage furrow ingression occurs between the two flagella. (G) At the end of the cell cycle, two daughter cells are formed, and each cell inherits a single kinetoplast, nucleus, basal body and flagellum. This figure was reproduced and modified from (Akiyoshi and Gull, 2013) under CC-BY.

It has been proposed that kinetoplastids might be among the earliest-branching eukaryotes based on various distinct features, such as the unique cytochrome c biogenesis machinery and unconventional kinetochore components (Allen et al., 2008; Cavalier-Smith, 2010; Akiyoshi, 2016). Understanding the biology of *T. brucei* could therefore provide insights into the origin of the spindle checkpoint and/or regulatory mechanisms for organelle segregation. I previously showed in procyclic (insect) form trypanosome cells that TbMad2 localizes at the basal body area throughout the cell cycle (Akiyoshi and Gull, 2013), which was verified in other studies (Dean et al., 2017; Zhou et al., 2019). No kinetochore signal was observed even when spindle microtubules were disrupted (Akiyoshi and Gull, 2013; Zhou et al., 2019), suggesting that TbMad2 unlikely plays a role in the canonical spindle checkpoint. In this study, I immunoprecipitated TbMad2 and identified another protein that localizes at the basal body area. I also identified two proteins that transiently localize at the basal body area.

## Results and Speculations

### Identification of a TbMad2-binding protein, TbMBP65

Mad2 belongs to the HORMA domain family, which includes Hop1, Rev7, and Mad2 (Aravind and Koonin, 1998; Rosenberg and Corbett, 2015; Tromer et al., 2019; Ye et al., 2020). Multiple sequence alignment of Mad2 proteins from various eukaryotes clearly shows that TbMad2 has high higher sequence similarity to Mad2 than to other HORMA domain containing proteins (Figure S1 and data not shown). Furthermore, many residues that are important for Mad2’s functions (Luo et al., 2000, 2002; Sironi et al., 2002; De Antoni et al., 2005; Mapelli et al., 2007) are conserved in TbMad2 (Figure S1). However, TbMad2 localizes at the basal body region (Akiyoshi and Gull, 2013; Dean et al., 2017; Zhou et al., 2019). To identify its interaction partners, I immunoprecipitated the YFP-tagged TbMad2 protein from procyclic cells and performed mass spectrometry (Table S1). This led to the identification of Tb927.6.1220 (Figure 2A), which was recently identified in the BioID of TbSpef1 (Sperm flagellar protein 1: also known as CaLponin-homology And Microtubule-associated Protein (CLAMP) or TbCMF18) (Chan et al., 2005; Dougherty et al., 2005; Broadhead et al., 2006; Baron et al., 2007; Gheiratmand et al., 2013; Dong et al., 2020). TbSpef1 localizes at the microtubule quartet that originates from the basal body and extends toward the anterior end of the cell. As previously shown (Dong et al., 2020), Tb927.6.1220 localizes at the basal body area throughout the cell cycle (Figure 2B). Some cells additionally had signals near the anterior tip (Figure 2B, arrowhead). Interestingly, I found a putative Mad2-interacting motif (Figure 2C), suggesting that Tb927.6.1220 may directly interact with TbMad2. I therefore propose to call this protein TbMBP65 for Tb<u>M</u>ad2-<u>B</u>inding <u>P</u>rotein <u>65</u> kDa. Its putative homologs were identified in kinetoplastids, diplonemids, and euglenids (Figure S2), but not outside of Euglenozoan species. The putative Mad2-interacting motif of TbMBP65 homologs is present in most kinetoplastids but is largely absent in diplonemids and euglenids. It is possible that TbMBP65 is a receptor at the basal body area that recruits TbMad2, similarly to how Mad1 recruits Mad2 onto kinetochores in other eukaryotes. Alternatively, TbMBP65 might play an effector role, similarly to the function of Cdc20 in the canonical spindle checkpoint.

**Figure 2.**
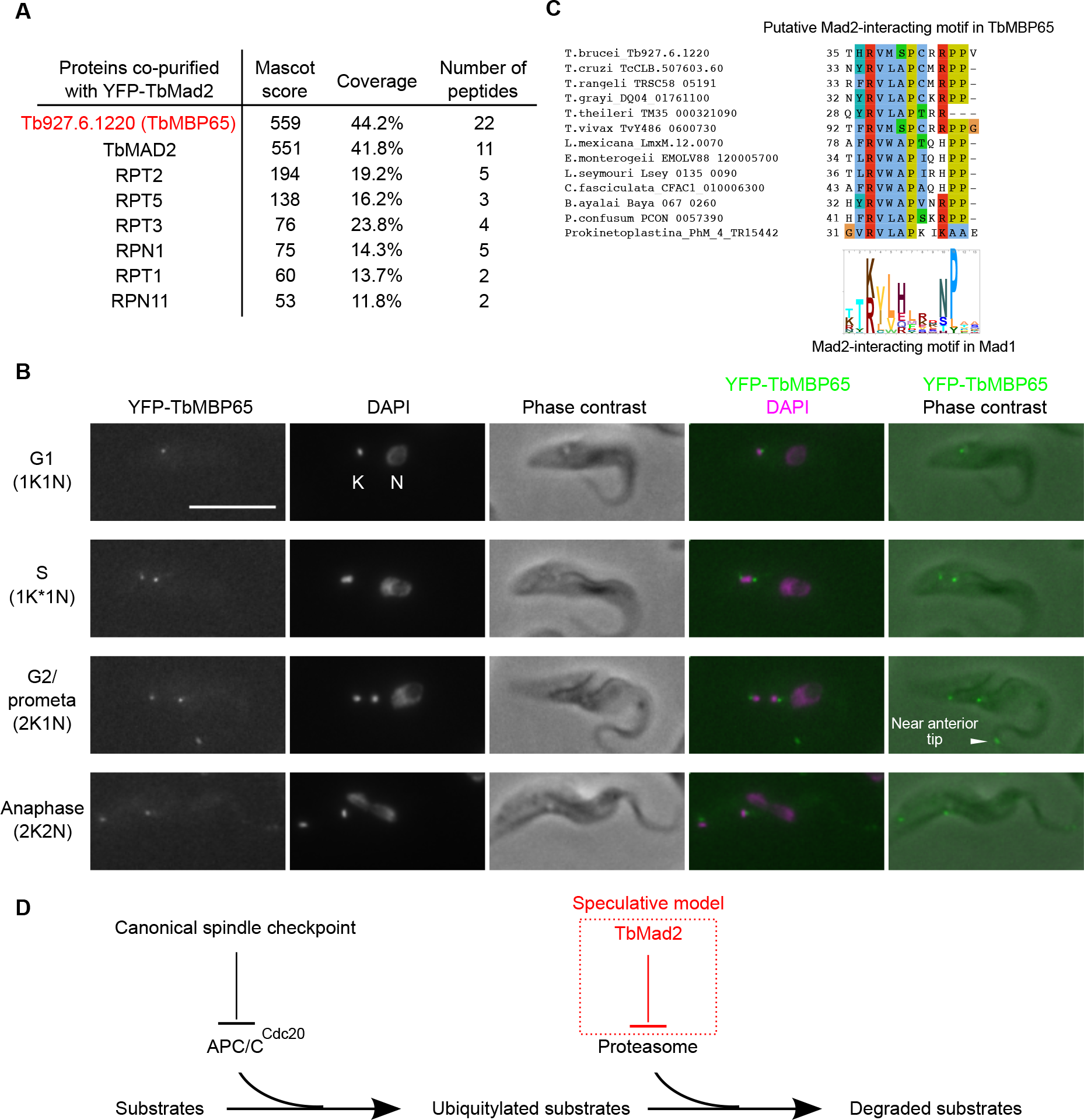
Identification of a putative TbMad2-binding protein, TbMBP65. (A) Summary of proteins detected in the immunoprecipitate of YFP-TbMad2 by mass spectrometry. See Table S1 for all proteins identified by mass spectrometry. (B) YFP-TbMBP65 localizes near basal bodies throughout the cell cycle. Note that some cells additionally have signals near the anterior tip (arrowhead). One allele of TbMBP65 was endogenously tagged at the N-terminus with YFP. Cells were fixed with 4% formaldehyde and stained with DAPI. Scale bar, 10 µm. (C) Alignment of TbMBP65 homologs in kinetoplastids reveals a putative Mad2-interacting motif. Logos of the Mad2-interacting motif in Mad1 proteins are shown below. (D) A highly speculative hypothesis that TbMad2 might negatively regulate proteasome activity, which would differ from the canonical spindle checkpoint that regulate APC/C^Cdc20^ activity.

To my surprise, TbMad2 also co-purified with RPT1/2/3/5 and RPN1/11 (Figure 2A), which are subunits of the 19S regulatory particle of the 26S proteasome (Li and Wang, 2002; Bard et al., 2018). These proteins were not detected in the immunoprecipitate of any other protein I purified using the same method (Table S1 and data not shown), suggesting that the observed co-purification is likely significant. Although totally speculative, it is interesting to discuss a possibility that TbMad2 might regulate the activity of proteasomes, which would differ from the canonical spindle checkpoint that more indirectly prevents substrate degradation via inhibition of the APC/C ubiquitin ligase (Figure 2D). Given that kinetoplastids might be one of the earliest-branching eukaryotes (Cavalier-Smith, 2010; Akiyoshi, 2016), the interaction between TbMad2 and proteasome regulatory subunits might represent an ancient mode for the regulation of the ubiquitin/proteasome-dependent substrate degradation system. It is noteworthy that another divergent eukaryote *Giardia intestinalis* (Metamonads) lacks APC/C components but has a Mad2 homolog (Eme et al., 2011; Gourguechon et al., 2013; Vicente and Cande, 2014; Markova et al., 2016). It will be interesting to examine whether GiMad2 co-purifies with proteasome subunits.

It is thought that basal bodies/flagella (centrioles/cilia) were already present in the last eukaryotic common ancestor (Carvalho-Santos et al., 2011), so it is conceivable that transmission of these organelles was critical for survival and that early eukaryotes might have had a checkpoint to monitor their proper inheritance. Given that TbMad2 localizes at the basal body area, it might regulate and/or monitor the duplication or segregation of basal bodies (or associated structures) in *T. brucei*, which possibly represents a prototype of the spindle checkpoint.

### TbAUK3 and its putative binding protein TbABP79 transiently localize at the basal body area

Aurora kinases play various key roles in eukaryotes, including regulation of mitosis and flagellar/cilia disassembly (Pan et al., 2004; Marumoto et al., 2005; Pugacheva et al., 2007; Inoko et al., 2012; Hochegger et al., 2013; Carmena et al., 2015; Korobeynikov et al., 2017; Bertolin and Tramier, 2020). *Trypanosoma brucei* has three Aurora kinases, TbAUK1, TbAUK2, and TbAUK3, among which TbAUK1 has a similar localization pattern as the Aurora B kinase in human (Li et al., 2008). As part of an effort towards identifying more mitotic proteins, I examined the localization of TbAUK2 and TbAUK3 (Tu et al., 2006; Jones et al., 2014; Fernandez-Cortes et al., 2017; Stortz et al., 2017) (Figure S3). TbAUK2 had a diffuse nuclear signal (data not shown), while TbAUK3 localized at the basal body area in a cell cycle dependent manner (Figure 3). Interestingly, TbAUK3 signal was not found in all 1K1N cells, suggesting that TbAUK3 starts to localize in late G1. In 1K*1N cells, there were multiple signals, which may reflect its localization at duplicated basal bodes or associated structures. Then the signals disappeared once the kDNA appeared as two distinct dots.

**Figure 3.**
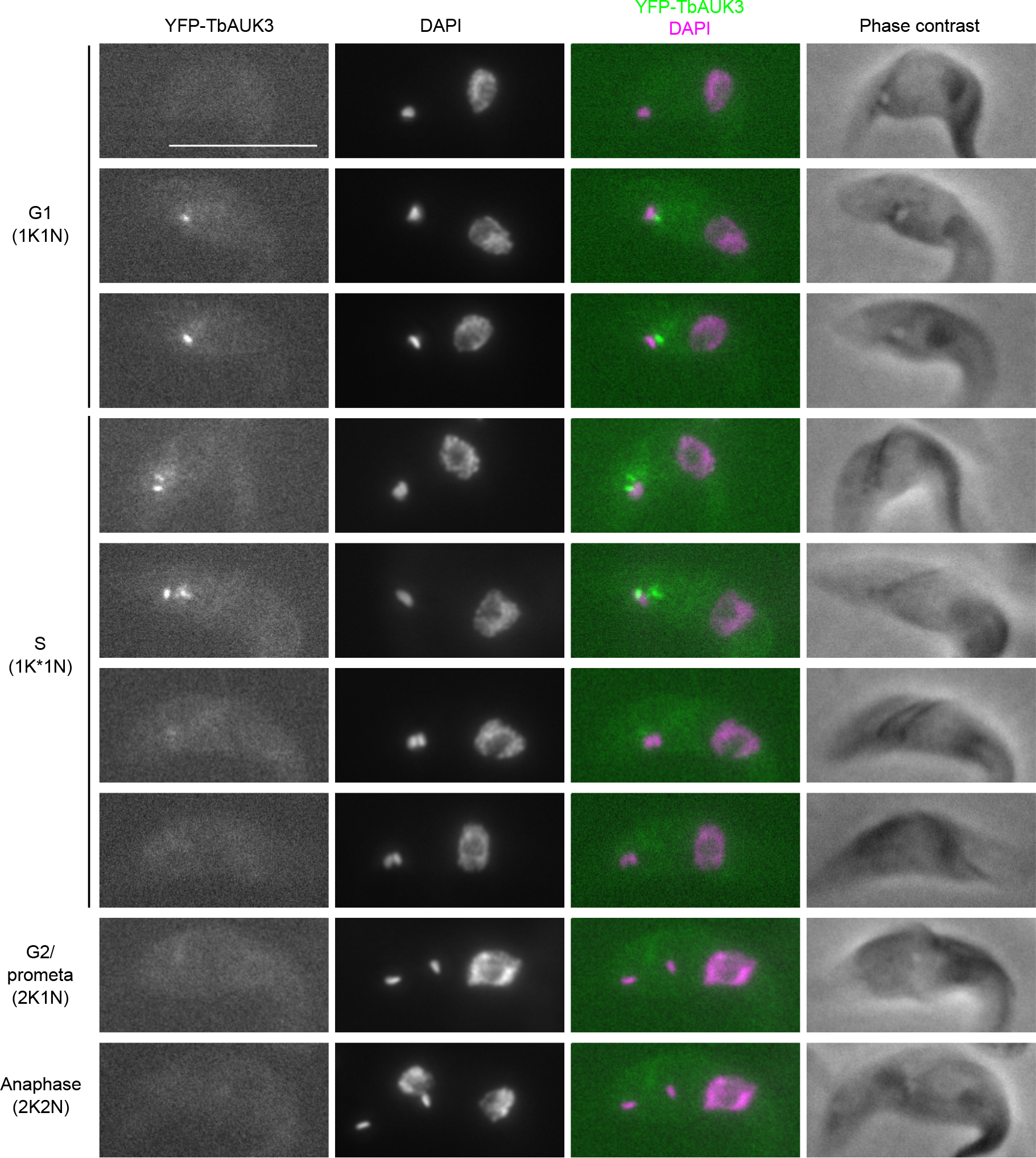
TbAUK3 localizes transiently at the basal body area. One allele of TbAUK3 was endogenously tagged at the N-terminus with YFP. Cells were fixed with 4% formaldehyde and stained with DAPI. Scale bar, 10 µm.

I then performed its immunoprecipitation and mass spectrometry, identifying Tb927.11.340 as a hit (Figure 4A and Table S1). This protein had a largely similar localization pattern as TbAUK3 (Figure 4B), so I propose to call it TbABP79 for Tb<u>A</u>UK3-<u>B</u>inding <u>P</u>rotein <u>79</u> kDa. No obvious homolog was found outside of kinetoplastids (Figure S4). The unique localization pattern of TbAUK3 and TbABP79 raises a possibility that these proteins might be involved in the biogenesis or separation of basal bodies or associated structures. Alternatively, they might monitor the segregation status of basal bodies or kinetoplasts.

**Figure 4.**
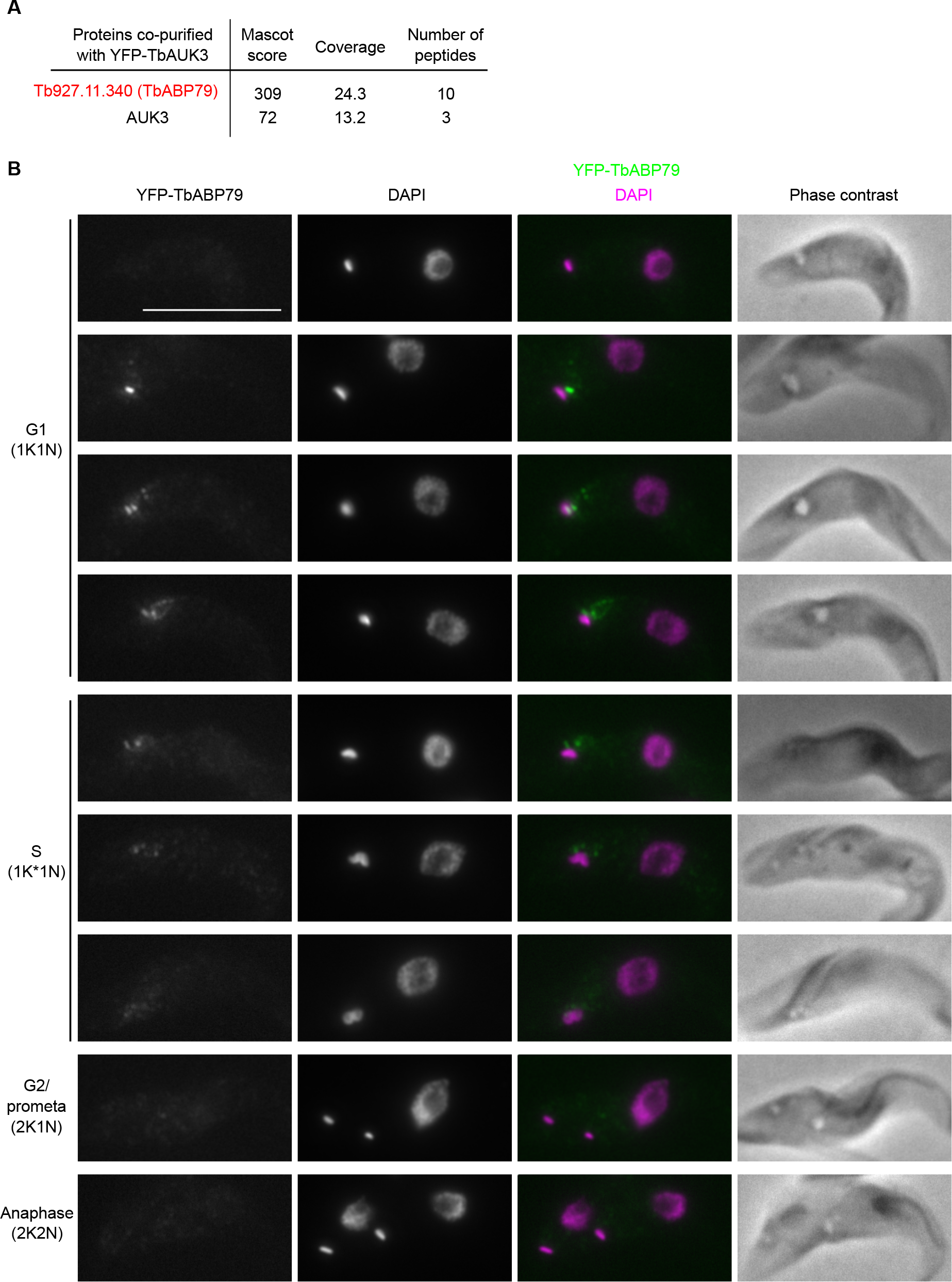
A putative TbAUK3-binding protein, TbABP79, also localizes transiently at the basal body area. (A) Summary of proteins detected in the immunoprecipitate of YFP-TbAUK3 by mass spectrometry. See Table S1 for all proteins identified by mass spectrometry. (B) One allele of TbABP79 was endogenously tagged at the N-terminus with YFP. Cells were fixed with 4% formaldehyde and stained with DAPI. Scale bar, 10 µm.

## Conclusions

In this manuscript, I reported several proteins that localize near basal bodies in *T. brucei*. I hope that this preprint will encourage some researchers to study these proteins in more detail and reveal their function in the future. It will be important to determine their exact location, for example by comparing with components of basal bodies, the microtubule quartet, hook complex components, and flagellar pocket collar (Lacomble et al., 2009; Esson et al., 2012; Albisetti et al., 2017; Halliday et al., 2019; Dong et al., 2020). It is also critical to reveal their knockdown or knockout phenotype. Some spindle checkpoint mutants show defects only in the presence of microtubule drugs (Hoyt et al., 1991; Li and Murray, 1991), so it might be necessary to identify proper stress conditions to detect a phenotype. I emphasize that there is currently no evidence to suggest that TbMad2 and TbAUK3 have similar functions. However, there are relationships among the ubiquitin-proteasome system, Aurora kinase, and basal body/centriole/cilia (Kasahara et al., 2014; Izawa et al., 2015; Shearer and Saunders, 2016), so it will be worth examining whether inhibition of TbMad2 or proteasome activity affects TbAUK3 and vice versa. Interestingly, treatment of procyclic trypanosome cells with a proteasome inhibitor lactacystin caused accumulation of cells with one kinetoplast and one nucleus with replicated DNA (Mutomba et al., 1997). Although lack of nuclear division can be explained by stabilization of cyclin B (Hayashi and Akiyoshi, 2018), why did these cells (apparently) have one kinetoplast and fail to divide? Because kinetoplast segregation depends on basal body separation (Robinson and Gull, 1991), one possible explanation is that proteasome activity may be required for the separation of basal bodies. In this scenario, the lack of cytokinesis could be explained by the hypothesis that initiation of cytokinesis may be linked to the completion of the kinetoplast segregation or basal body segregation in trypanosomes (Ploubidou et al., 1999). It will be interesting to revisit the phenotype of lactacystin-treated cells by examining the status of basal bodies, other associated structures, and cytokinesis.

## Materials and Methods

### Trypanosome cells and plasmids and microscopy

Trypanosome cell lines, plasmids, and primers used in this study are listed in Table S2, S3, and S4. All cell lines used in this study were derived from *T. brucei* SmOxP927 procyclic form cells (TREU 927/4 expressing T7 RNA polymerase and the tetracycline repressor to allow inducible expression) (Poon et al., 2012). Cells were grown at 28°C in SDM-79 medium supplemented with 10% (v/v) heat-inactivated fetal calf serum (Brun and Schönenberger, 1979). Endogenous YFP tagging constructs were made using the pEnT5-Y vector (Kelly et al., 2007) with the DNA Strider software (Marck, 1988). Transfected cells were selected by the addition of 25 µg/ml hygromycin. Fluorescence microscopy was performed as described previously (Akiyoshi and Gull, 2014).

### Immunoprecipitation and mass spectrometry

Immunoprecipitation of YFP-TbMad2 and YFP-TbAUK3 was carried out using anti-GFP antibodies as described previously (Akiyoshi and Gull, 2014) with the following modifications. 400 ml cultures of asynchronously growing cells were harvested at ∼1.2 x 10^7^ cells/ml. Cells were pelleted by centrifugation (900 g, 15 min), washed once with PBS, resuspended in 0.5 ml of BH0.15 (25 mM HEPES pH 8.0, 2 mM MgCl_2_, 0.1 mM EDTA, 0.5 mM EGTA, 1% NP-40, 150 mM KCl, and 15% glycerol) with protease inhibitors (20 µg/ml each of leupeptin, pepstatin, and E-64 as well as 0.2 mM PMSF) and phosphatase inhibitors (1 mM sodium pyrophosphate, 2 mM Na-beta-glycerophosphate, 0.1 mM Na_3_VO_4_, 5 mM NaF, and 200 nM microcystin-LR), and then frozen in small drops into liquid nitrogen in a mortar. Cells were ground with a pestle with liquid nitrogen, and the cell power was transferred into a 50 ml tube. When the samples started to melt, 1 ml of BH0.15 with inhibitors was added, followed by 25 min incubation to allow lysis on ice. The samples were then centrifuged at 14,000 rpm for 30 min at 4 °C. The supernatant was transferred into a new tube and was mixed with mouse monoclonal anti-GFP antibodies (11814460001, Roche) that had been pre-conjugated with Protein-G Dynabeads with dimethyl pimelimidate using a protocol described in (Unnikrishnan et al., 2012). Subsequent steps and mass spectrometry were performed essentially as described (Akiyoshi and Gull, 2014), and peptides were identified with MASCOT (Matrix Science). Proteins identified with at least two peptides were considered as significant and shown in Table S1.

### Sequence analysis

Protein sequences and accession numbers for proteins used in this study were retrieved from TriTryp database (Aslett et al., 2010), Wellcome Sanger Institute (https://www.sanger.ac.uk/), UniProt (UniProt Consortium, 2019), or published studies (O’Neill et al., 2015; Butenko et al., 2020). Homology searches were done using BLAST in TriTryp (Aslett et al., 2010), HMMER web server (Potter et al., 2018), or hmmsearch using manually prepared HMM profiles (HMMER version 3.0) (Eddy, 1998). Multiple sequence alignment was performed with MAFFT (L-INS-i method, version 7) (Katoh et al., 2019) and visualized with the Clustalx coloring scheme in Jalview (version 2.11) (Waterhouse et al., 2009). The HMM logo for the Mad2-interacting motif in Mad1 was created with skylign (Wheeler et al., 2014) based on Mad1 homologs from various eukaryotes (van Hooff et al., 2017). Coiled coils were predicted using COILS program (Lupas et al., 1991).

## Acknowledgments

I thank Professor Keith Gull for his support. I also thank the Advanced Proteomics Facility, Benjamin Thomas, and Gabriela Ridlova for mass spectrometry analysis. Experiments reported in this manuscript were carried out when I was supported by postdoctoral fellowships from the EMBO and Human Frontier Science Program. I am currently supported by a Wellcome Trust Senior Research Fellowship (grant no. 210622/Z/18/Z) and the European Molecular Biology Organization Young Investigator Program. I declare that no competing interests exist.

## Supplemental Materials

### Supplemental Figures

**Figure S1.**
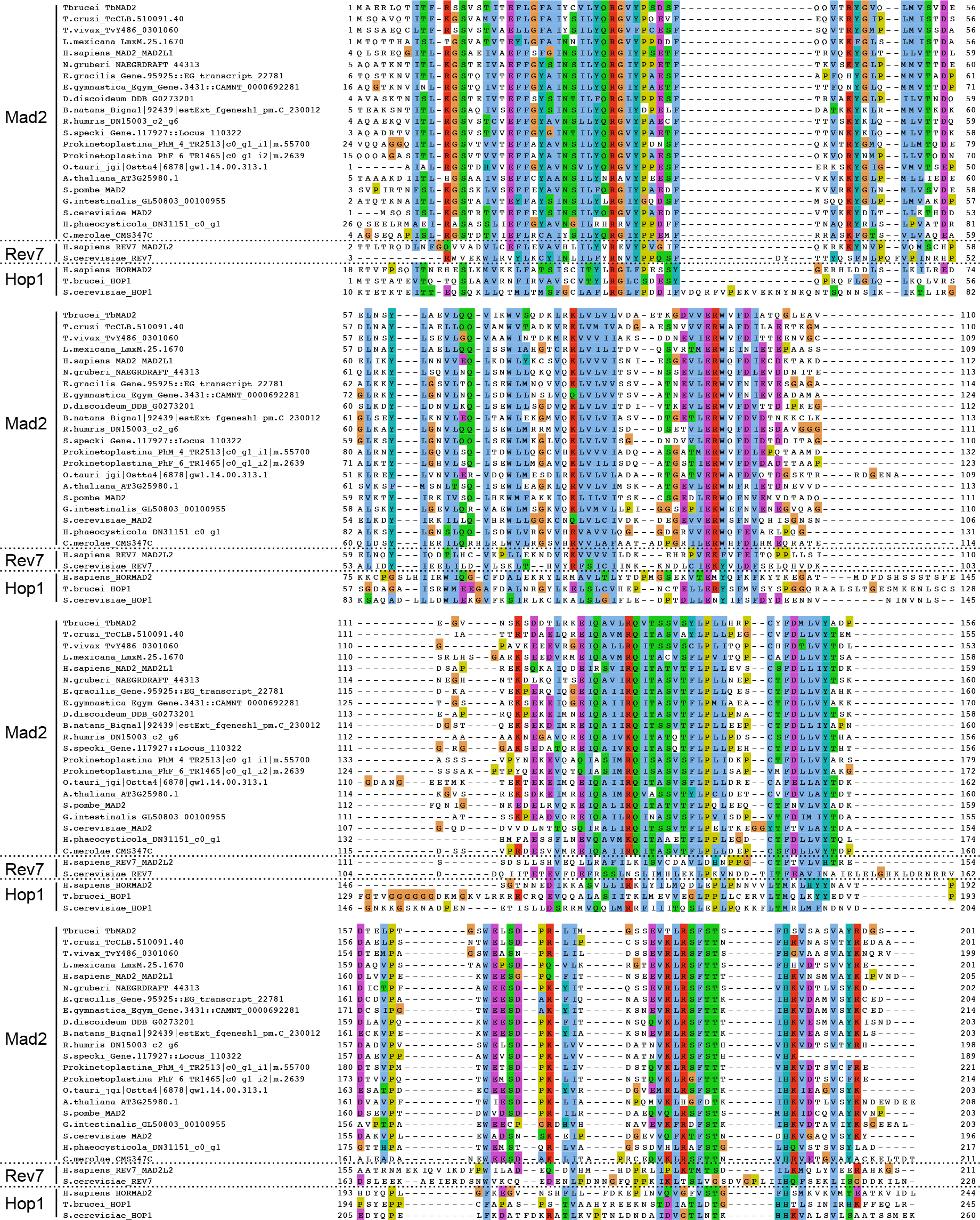
Sequence analysis of TbMad2. Multiple sequence alignment of Mad2 from various eukaryotes as well as Rev7 and Hop1 from some species, showing that TbMad2 has higher sequence similarity to Mad2 than to other HORMA domaining containing proteins.

**Figure S2.**
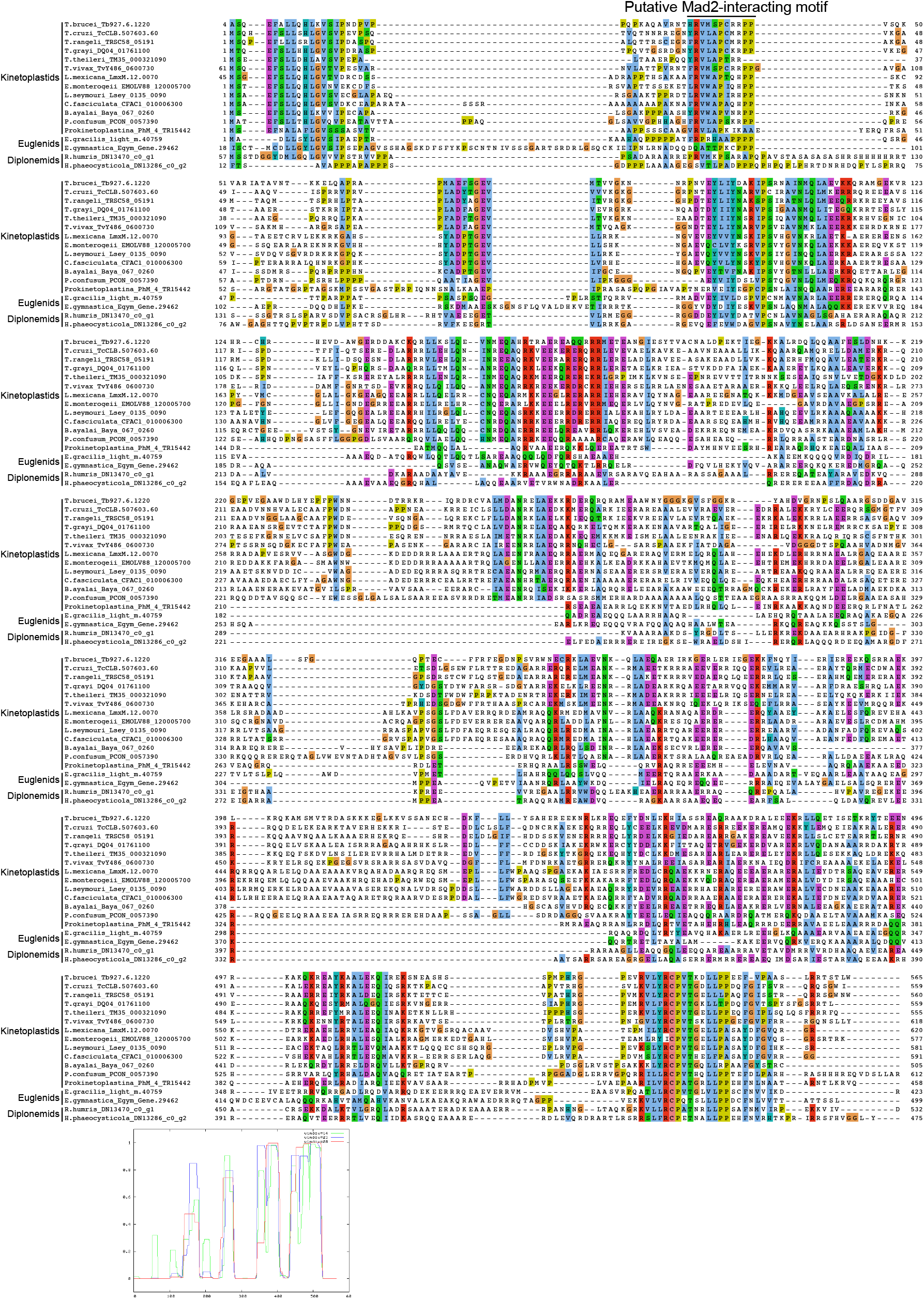
Sequence analysis of TbMBP65. Multiple sequence alignment of TbMBP65. Its putative homologs are found only in Euglenozoa species. A putative Mad2-binding motif and coiled coil prediction for TbMBP65 are shown.

**Figure S3.**
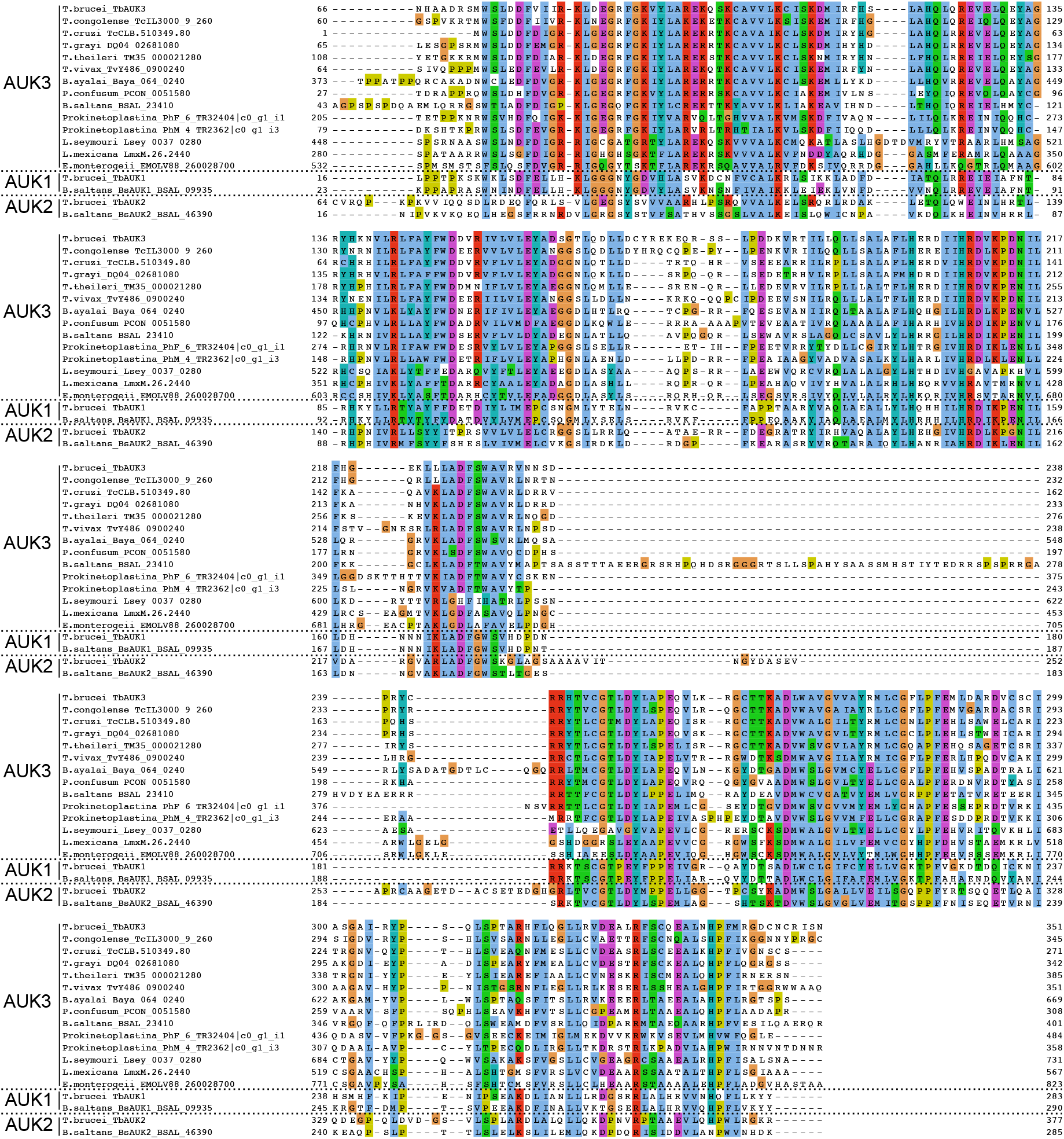
Comparison of the kinase domain of AUK1, AUK2, and AUK3 in kinetoplastids. Multiple sequence alignment of the kinase domain of Aurora kinases in kinetoplastids with emphasis on AUK3. Putative AUK3 homologs are found even in some Prokinetoplastina.

**Figure S4.**
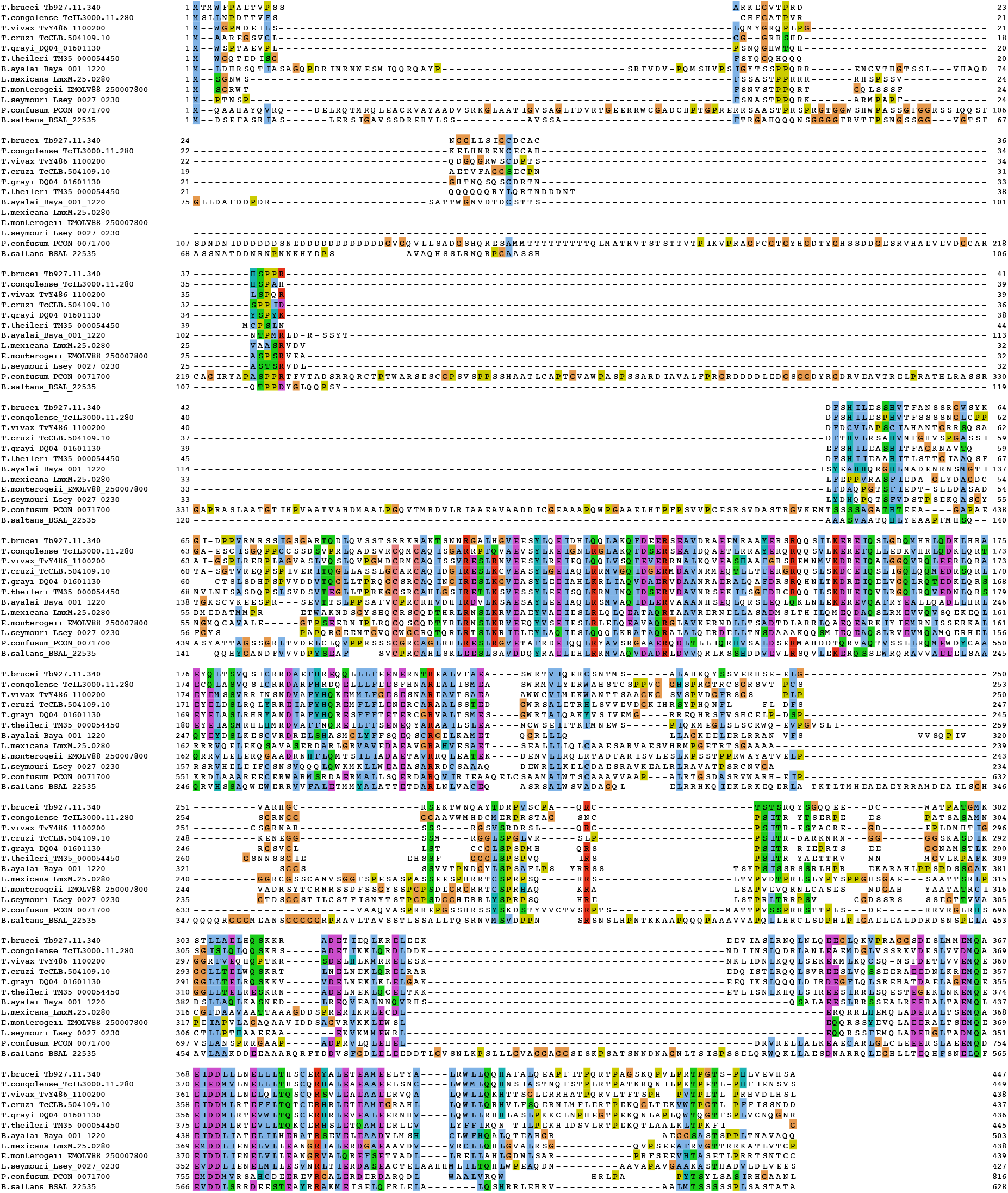
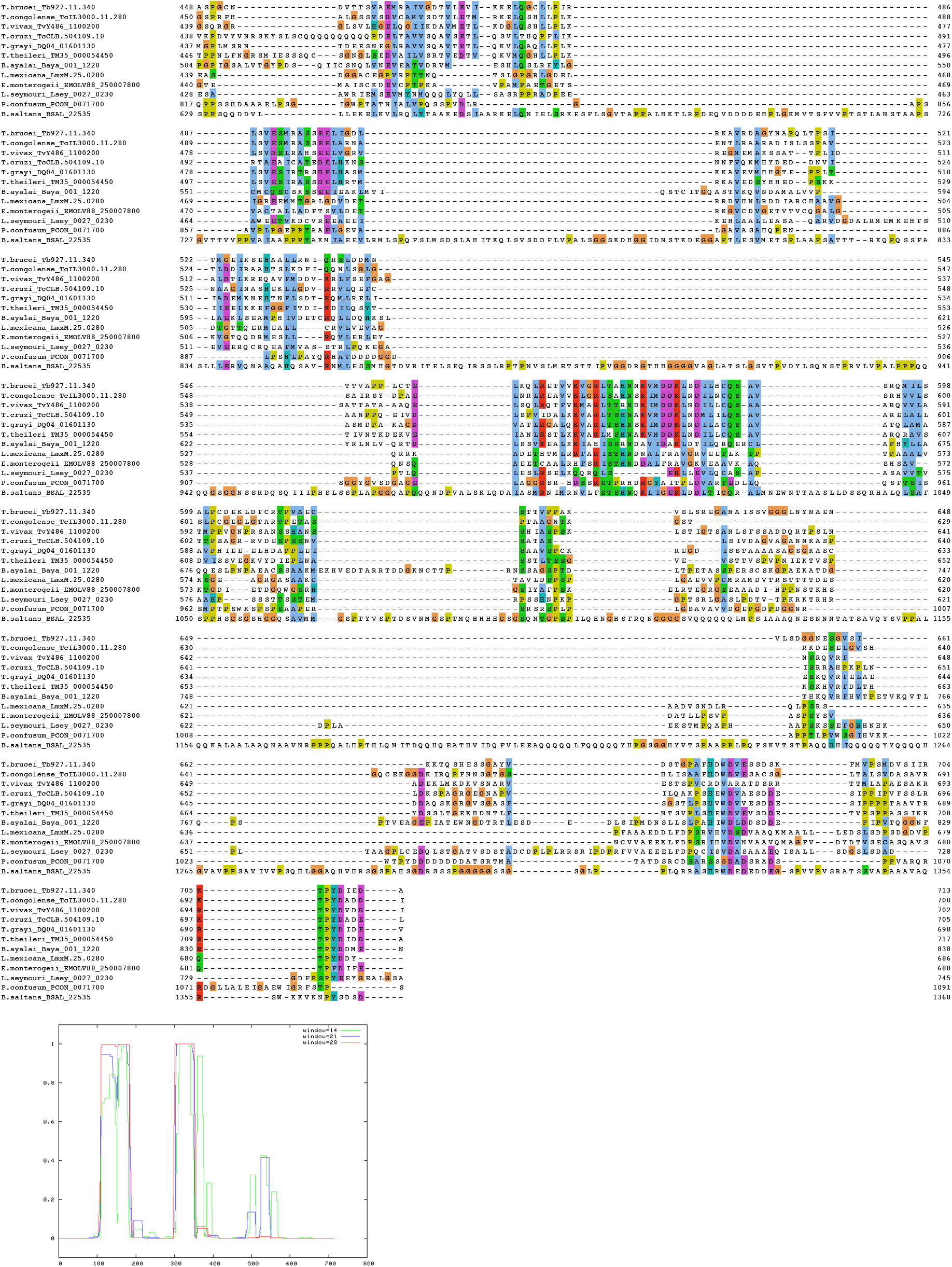
Sequence analysis of TbABP79. Multiple sequence alignment of TbABP79 homologs in kinetoplastids.

### Supplemental Tables

**Table S1: List of proteins identified in the immunoprecipitates of YFP-TbMad2 and YFP-TbAUK3 by mass spectrometry (Excel file)**

**Table S2.** Trypanosome cell lines used in this study.

| Name | Description |
| --- | --- |
| SmOxP9 | Parental cell line, TREU 927/4 procyclic form cells expressing that expresses TetR and T7 RNAP (Poon et al., 2012) |
| BAP72 | YFP-TbMad2 (Akiyoshi and Gull, 2013) |
| BAP59 | YFP-TbMBP65 (this study) |
| BAP76 | YFP-TbAUK3 (this study) |
| BAP81 | YFP-TbABP79 (this study) |

**Table S3.** Plasmids used in this study.

| Name | Description |
| --- | --- |
| pEnT5-Y | TY-YFP tagging vector, Hygromycin (Kelly et al., 2007) |
| pBA2 | TY-YFP-TbMad2 tagging construct, Hygromycin (this study) |
| pBA38 | TY-YFP-TbMBP65 tagging construct, Hygromycin (this study) |
| pBA12 | TY-YFP-TbAUK3 tagging construct, Hygromycin (this study) |
| pBA54 | TY-YFP-TbABP79 tagging construct, Hygromycin (this study) |

**Table S4.**
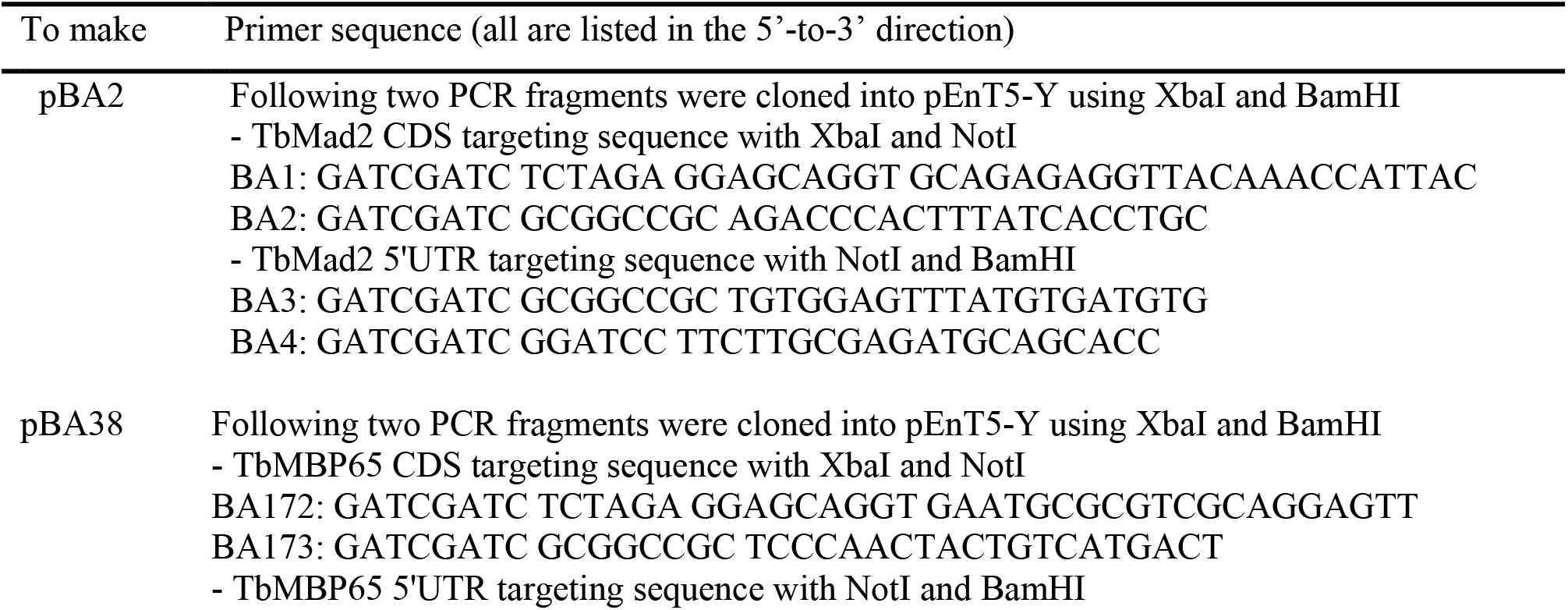
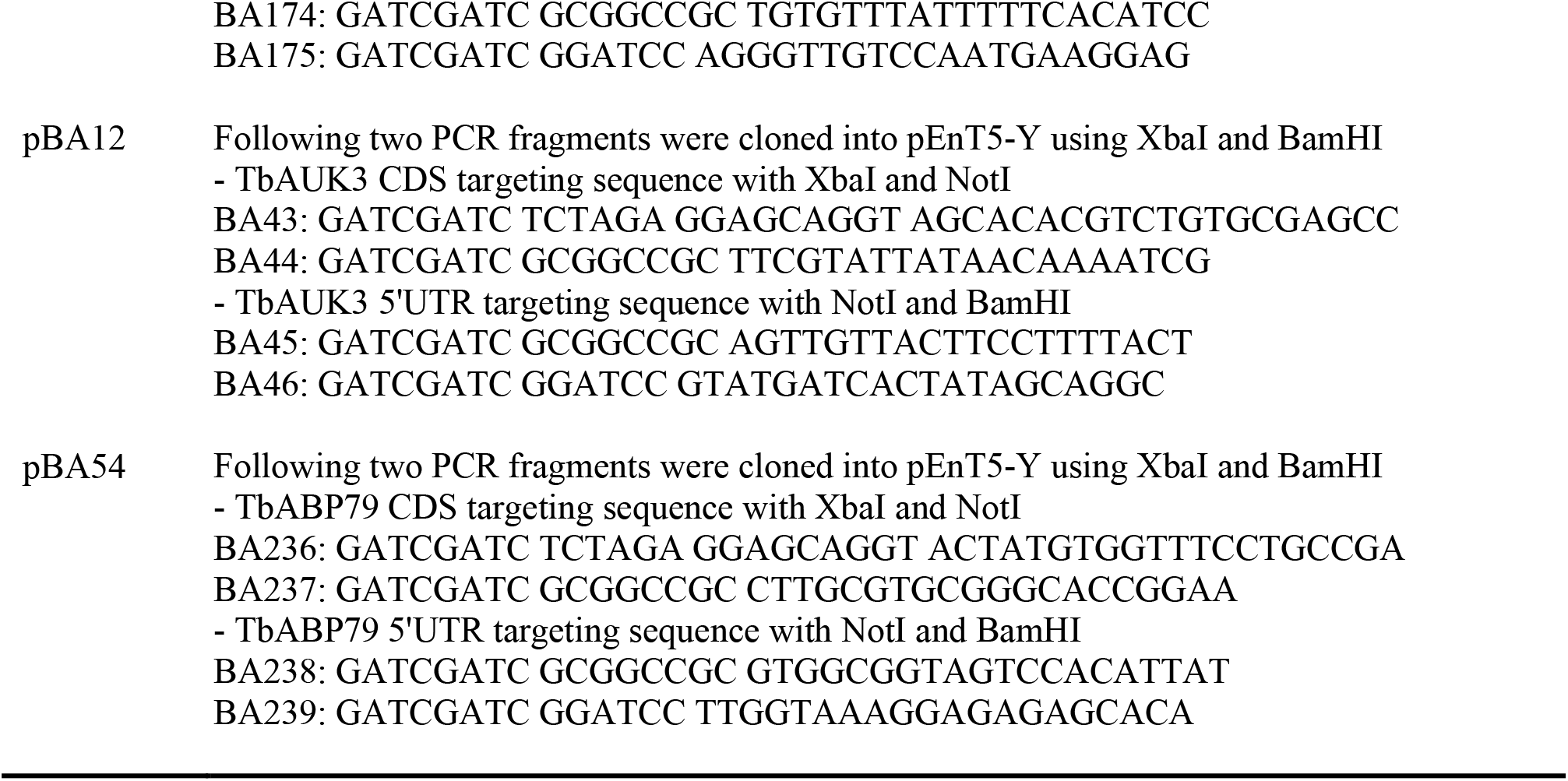
Primers used in this study.

